# Automated Change Detection Methods for Satellite Data that can Improve Conservation Implementation

**DOI:** 10.1101/611459

**Authors:** Michael J. Evans, Jacob W. Malcom

## Abstract

A significant limitation in biodiversity conservation has been the effective implementation of laws and regulations that protect species habitats from degradation. Flexible, efficient, and effective monitoring and enforcement methods are needed to help conservation policies realize their full benefit. As remote sensing data become more numerous and accessible, they can be used to identify and quantify land cover changes and habitat loss. However, these data remain underused for systematic conservation monitoring in part because of a lack of simple tools. We adapted and developed two generalized methods that automatically detect land cover changes in a variety of habitat types using free and publicly available data and tools. We evaluated the performance of these algorithms in two ways. First, we tested the algorithms over 100 sites of known change in the United States, finding these approaches were effective (AUC > 0.90) at distinguishing between areas of land cover change and areas of no change. Second, we evaluated algorithm effectiveness by comparing results to manually identified areas of change in four case studies involving imperiled species habitat: oil and gas development in the range of the Greater Sage Grouse; sand mining operations in the range of the dunes sagebrush lizard; loss of Piping Plover coastal habitat in the wake of hurricane Michael (2018); and residential development in beach mouse habitat. The relative performance of each algorithm differed in each habitat type, but both provided effective means of detecting and delineating habitat loss. Our results show how these algorithms can be used to help close the implementation gap of monitoring and enforcement in biodiversity conservation and we provide a free online tool that can be used to run these analyses.

**Article impact statement:** Methods for automating the detection of habitat loss in satellite images that can be used to monitor and enforce conservation policy.

## Introduction

Biodiversity around the world is threatened with annihilation (Ceballos et al. 2017), and direct habitat loss is the leading cause of the loss of individuals and species extinctions (Newbold et al. 2015; Betts et al. 2017). To address this threat, biodiversity conservation is often focused on protecting species’ habitat through a variety of legal and policy mechanisms (UN Environment World Conservation Monitoring Centre & International Union for Conservation of Nature 2017). However, there is reason to believe entities with regulatory authority may lack the means to monitor and enforce protections. Without regular monitoring and enforcement, conservation laws may be nothing more than paper tigers (Salomon et al. 2014). Options for monitoring and enforcing laws that protect habitat have historically required time intensive efforts on the ground, and thus have been limited by funding and personnel availability. However, technological advances are expanding the options for cost effective monitoring efforts (e.g., aquatic telemetry; Hussey et al. 2015, remote cameras detecting poachers; Hossain et al. 2016). Given the central role that habitat conservation plays in conserving imperiled species, methods to automatically detect habitat loss in near real-time could significantly enhance compliance monitoring and enforcement capabilities, and substantially increase the effectiveness of conservation laws.

Many conservation laws include provisions to protect habitat. For example, the U.S. Endangered Species Act (ESA) is the primary tool for conserving imperiled species in the United States. Among its strengths are the requirement for identification of ‘critical habitat’ that is necessary for the conservation of listed species, and prohibitions against destroying or adversely modifying these habitats (United States Congress 1978). Similarly, Japan’s Nature Conservation Law designates ‘Wilderness’ and ‘Nature Conservation’ areas (Japan National Diet 1972); the New Zealand Conservation Act created several specially protected areas (New Zealand Parliament 1987.); and various international agreements include provisions to reduce habitat loss (e.g. Convention on Biological Diversity; UN Sustainable Development Goals). As written, these laws, policies, and treaties should be stopping or significantly slowing habitat loss and degradation. But this conclusion depends heavily on a critical assumption: that these laws are implemented as written, including monitoring of conservation agreements and enforcement of prohibitions. That assumption is often not independently tested, and the continued loss of species and their habitats indicate there is a substantial implementation gap (López-Bao et al. 2015; Chapron 2017).

Enforcement is a critical component of any law. Without compliance monitoring and punishment of infractions there is little reason to think legal protections will be effective at changing outcomes (Keane et al. 2008; Trouwborst et al. 2017). For example, if drivers believe there is little risk of punishment for exceeding a speed limit because there is no monitoring, there is every reason to believe they will. Currently, there is little research available on the extent of enforcement and compliance of habitat protection laws and policies (Malcom et al. 2017), but there are many reasons to believe they are lacking. Staff at U.S. federal agencies have acknowledged that they lack the resources to carry out even basic compliance monitoring and are unable to read monitoring reports submitted by permittees, much less carry out independent monitoring (Government Accountability Office 2009). Furthermore, the two federal agencies responsible for implementing the ESA, the U.S. Fish and Wildlife Service and National Marine Fisheries Service (hereafter ‘the Services’), have finalized over a thousand conservation agreements, many of which authorize limited habitat destruction in exchange for mitigation to offset some of those effects (Malcom & Li 2015). Perennial funding shortages, however, have left the Services unable to monitor for compliance with many of those agreements; a shortcoming that threatens to undermine the potential benefits of the ESA. These basic examples of a lack of monitoring and enforcement highlight a critical weakness in the implementation of conservation law and consequently the protection of biodiversity.

Insufficient monitoring undermines imperiled species conservation in two ways. First, habitat protections may go unenforced. For example, satellite images revealed that under a habitat conservation plan for the eastern indigo snake (*Drymarchon couperi*) in Georgia, USA, over half of a forest parcel had been cleared despite the requirement that the permittee manage the parcel for the species until at least 2027 (Malcom 2017). Situations like this are a double blow for species: not only has authorized habitat loss occurred, but the conservation measures to minimize or offset those losses were never fully realized. Second, inadequate monitoring leaves conservationists in the dark about the status of species’ habitat. If 60 percent of a species’ habitat has been degraded, that knowledge should factor into decision making. Left unresolved, a lack of monitoring could negate the expensive and difficult work of securing legal protections for species and their habitats and negotiating conservation agreements.

Although the challenge of inadequate enforcement is not new, solutions to date have relied heavily on increased financial support for field work (Chandra & Idrisova 2011). This strategy may be untenable at broad scales given inconsistent and decreasing political will and concomitant funding declines (McCarthy et al. 2012; Waldron et al. 2013). Even when government agencies monitor for compliance with certain projects, they may lack the ability to do so regularly. Monitoring that occurs intermittently leaves ample opportunities for noncompliance in the interim. By the time violations are identified, the environmental damage may be irreversible. Large-scale monitoring programs to efficiently and automatically detect disturbances to wildlife habitat are needed.

The growing availability of free satellite images and other remote-sensing data provide an efficient and effective solution for many biodiversity monitoring challenges (Turner et al. 2003). When combined with information on species range and areas permitted for habitat disturbance or destruction, these data open a wealth of opportunities for compliance monitoring and enforcement. As remote sensing data has become more ubiquitous and accessible, so too have the number of approaches for change detection (Willis 2015). Often these analyses focus on one land cover type, with most of the research focused on forest loss (Potapov et al. 2008; Hansen et al. 2010; Song et al. 2018). A significant challenge now is to expand the generality of algorithms to automate change detection across habitats, which would enable and simplify monitoring at regional and continental scales.

Here we report on two automated land cover change-detection algorithms developed to aid conservation compliance monitoring across different habitat types and at broad scale. We evaluate the utility of these methods using systematically collected validation data and four case studies. Both algorithms use data that is readily available online, meaning anyone, including government agencies, conservation organizations, and the public, can use them to improve conservation. We demonstrate that these approaches are sufficiently effective, efficient, and flexible for use in large- and small-scale systematic conservation monitoring efforts. Adoption of automated change detection can help close one of the biggest gaps in biodiversity conservation, and we discuss the potential for future technological and regulatory development to further leverage their potential.

## Methods

We used the Google Earth Engine platform, which provides real-time access to terabytes of remote sensing data and the cloud computing capabilities to analyze them (Gorelick et al. 2017), to create processes to automatically detect changes in land cover between satellite images collected over two time periods. We analyzed images from the Sentinel-2 satellite system. Sentinel-2 is deployed and maintained by the European Space Agency, providing global coverage of 10-meter resolution imagery every 12 days. Sentinel-2 images contain 13 bands that record reflectance values in the visible, near infrared, short-wave infrared, and near ultraviolet spectra (Drusch et al. 2012).

The basic process (Fig. 1) involves the following steps:

**Figure 1.**
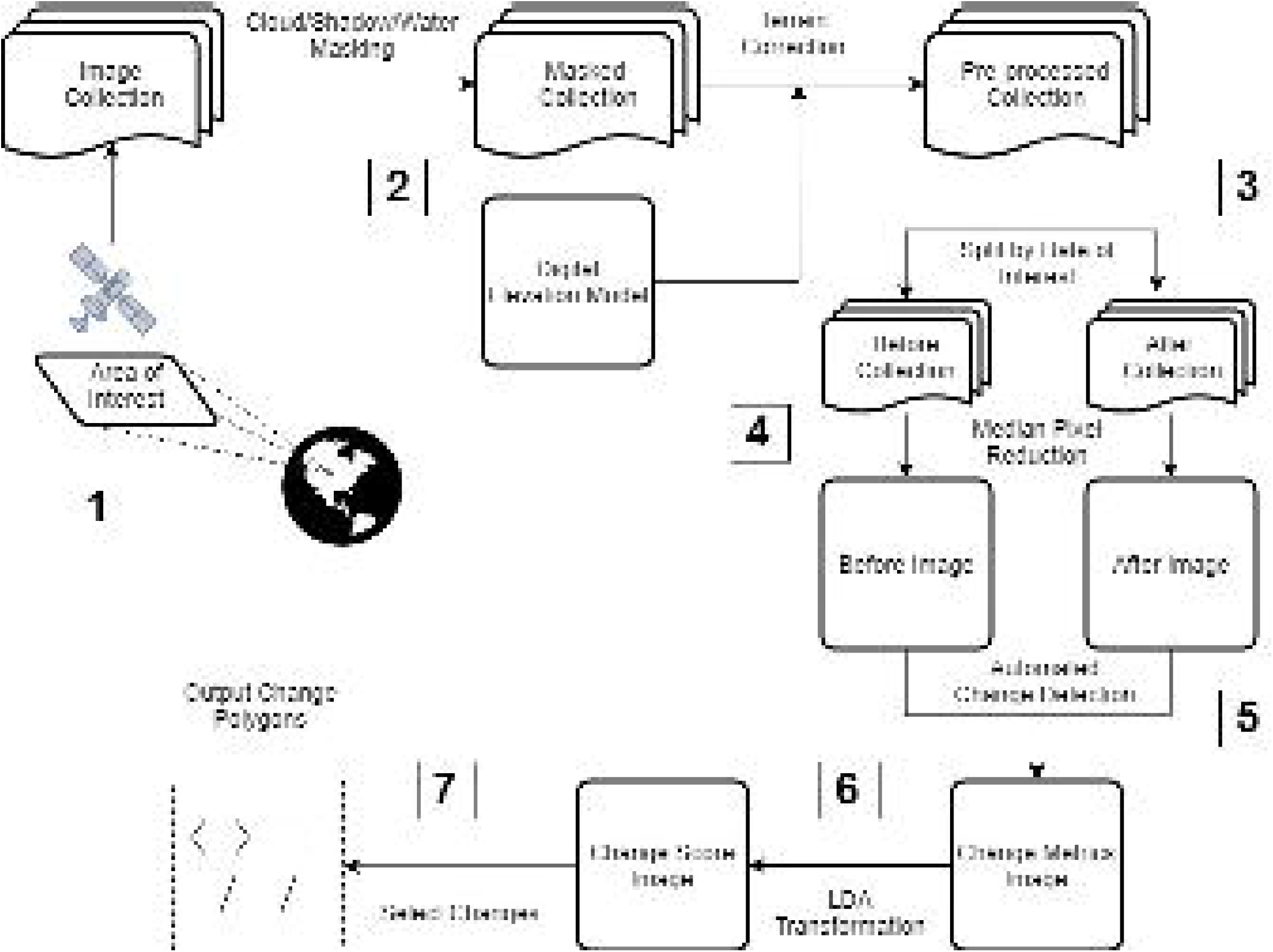
Conceptual model showing steps for processing and automatically detecting changes between two sets of satellite images used in this paper. Numbers correspond to the steps for selecting and preprocessing image collections and performing change detection between two time periods described in the text.

1. Define an area of interest and collect satellite images.
2. Process images (mask clouds, correct for terrain, etc.)
3. Divide images into before and after collections
4. Composite before and after collections into single images.
5. Calculate pixel-wise change metrics between images.
6. Identify minimum values that correspond to the change to be detected.
7. Select pixels exceeding these minimum change thresholds.

### Image Processing

After defining an area of interest and collecting all the spatially overlapping Sentinel-2 images, we first removed cloud and cloud shadow pixels from each image in the collection. Built-in cloud masking is limited for Sentinel-2 imagery, because this system does not contain a thermal sensor measuring temperature, which is critical to common cloud masking procedures (Zhu et al. 2015). We used the quality assurance bands included with all S2 images, and additionally calculated cloud and shadow probability metrics as follows.

To identify cloudy pixels, we implemented an adaptation of the simpleCloudScore algorithm developed for LandSat provided in Google Earth Engine, which uses a combination of indices to assign a per pixel cloud likelihood score from 0 to 1 (Supporting Information). We identified any pixels receiving a score of 0.15 or greater as cloud. We then calculated a set of likely cloud shadow locations by translating the location of cloud pixels in the *x* and *y* directions according to

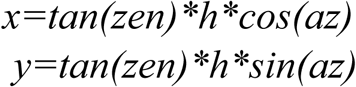

where *h* is the cloud height, *zen* and *az* are the sun zenith and azimuth at the time and location of the image, as recorded by Sentinel-2. This translation is applied using a set of possible cloud heights (*h*) to create a set of polygons encompassing possible cloud shadow locations (Zhu & Woodcock 2012). We then calculated the RGB ratio shadow indices (Sarabandi et al. 2004) for each pixel and classified those with a score over 0.25 as shadow (Supporting Information). Any of these shadow pixels that fell within the translated cloud locations were then labeled as cloud shadow. Finally, we identified water pixels using a set of water indices including a normalized difference water index (Xu 2006), and darkness indices to obtain a 0 -1 water likelihood score (Supporting Information). We masked any pixels over 0.25. The set of cloud, shadow, and water pixels were then removed from each image.

After all images in the spatially filtered collection were masked for clouds, shadows, and water, we applied per-pixel terrain correction using the *c-correction* equation (Teillet et al. 1982). This method standardizes the reflectance of sloped surfaces using the illuminance of each pixel as determined by a digital elevation model and the solar zenith and azimuth at the time and location of the image. We used the 30m resolution digital elevation model from the U.S. Geological Survey (Farr et al. 2007) to determine slope and aspect.

Change detection was ultimately run between single before and after images. However, clouds, shadows and other artefacts can make detection difficult between any two images. Therefore, we compared two composites of all the masked and corrected images in before and after time periods. By default, the before period contained a year of images preceding the date after which we wanted to check for changes, and a three-month period following this date comprised the after period. Following cloud/shadow/water masking and terrain correction, we created a single-image composite for each time period by selecting the median value of each pixel stack. These single before and after images were then clipped to the exact geometry of the study area and used as inputs to automated change detection algorithms. Six bands corresponding to blue, green, red, near infrared, short-wave infrared 1, and short-wave infrared 2 were used in all calculations of change.

### Change Detection Algorithms

While a variety of algorithms have been developed to detect changes between satellite images (Willis 2015) we started from two fundamentally different approaches. The first builds on the method used by the U.S Geological Survey to produce the National Land Cover Dataset land cover change (LCC) data (Jin et al. 2013) and uses features that relate to real phenomena. We refer to this as the LCC algorithm. First, six spectral change metrics are calculated between before and after imagery on a per-pixel basis:

1. The Change Vector (*CV*) measures the total change in reflectance values between two images across the visible and infrared spectrum.
2. Relative CV Maximum (*RCV*_*MAX*_) measures the total change in each band scaled to their global maxima.
3. Differences in Normalized Difference Vegetation Index (*dNDVI*) uses ratios between near infrared and red reflectance to indicate changes in the concentration of vegetation.
4. Ratio Normalized Difference Soil Index (*dRNDSI*) uses ratios between short-wave infrared and green reflectance to indicate changes in the concentration of bare ground.
5. Normalized Burn Ratio (*dNBR*) is the normalized difference between the green and short-wave infrared bands, indicating the severity of burned areas of vegetation.
6. Normalized Difference Water Index (*dNDWI*) is the normalized difference between the near- and short-wave-infrared bands, indicating moisture.

Calculating all six metrics at each pixel produces an unscaled change image with six bands (one band per metric). We then convert pixel values for each band to z-scores using the mean or minimum value and standard deviation of values across the image. We use global means for normalized indices (*dNDVI, dRNDSI, dNBR, dNDWI*), and global minimums for scaled indices (*CV* and *RCV*_*MAX*_) as in Jin et al. (2013). The output is a six-band image consisting of the standardized z-scores for each change metric. This transformation on the change image centers and scales per pixel changes relative to baseline changes in reflectance, brightness, etc. between the before and after images.

The output image is then iteratively re-weighted using the probability that a pixel represents no-change. We approximate this probability with *p-values* from the relevant distributions for each band. For normalized indices (*dNBR, dNDSI, dNDVI, dNDWI*) we use the cumulative distribution function of a standard normal distribution ∼*N(0, 1)*. The *CV* statistic is calculated as the sum of squared deviations for each image band, and therefore is approximately chi-square distributed (Lancaster & Seneta 2005). We calculated *p-values* using the cumulative distribution function of a chi-square distribution with degrees of freedom equal to the number of image bands minus one.

The second algorithm is the multivariate alteration detection (MAD) algorithm (Nielsen 2007). This approach uses canonical correspondence analysis to identify linear transformations that maximize correlation between two sets of variables, in this case, the bands of two images. Extreme deviations are identified by calculating the sum of squared deviations from the mean of each canonical variate, relative to its variance. We implement the MAD algorithm by performing singular value decomposition on a correlation matrix of the bands of two images. Singular value decomposition produces two orthogonal vectors, ***U*** and ***V*** which equate to the coefficients for the CCA linear transformation. Singular value decomposition also produces a diagonal vector ***S***, equivalent to the correlation coefficients (*ρ*_*i*_) of canonical correlation. Canonical variates are then obtained as the difference between the bands of the first image transformed by ***U*** and the bands of the second image transformed by ***V***.

The output of a single iteration of the MAD algorithm is an image with *min(m, n)* bands corresponding to the canonical variates ***V***, a band containing the chi-square summary statistic at each pixel (*χ*^*2*^), and a band containing the corresponding *p-value* from a chi-square distribution with *min(m, n)* degrees of freedom. As with the LCC algorithm, we then recalculate the MAD variates using these *p-values* to weight the calculation of means and variances as in Nielsen (2007). This procedure is performed iteratively until the output image has stabilized, or a maximum of *k =* 30 iterations has been reached (Nielsen 2007). We use changes in the correlation coefficients between iterations, *c*_*k*_ = *abs*|*max(****S***_***k***_*) – max(****S***_***k-1***_)| to evaluate convergence of the reweighting algorithm by inferring stability when *c*_*k*_ < 0.001. All image processing, calculations and transformations are performed in Google Earth Engine (Supporting Information).

### Algorithm Validation

We collected algorithm output at 100 study sites across the continental United States that had been manually identified as undergoing habitat loss due to anthropogenic landscape modification since 2016 (Supporting Information). The predominant land cover undergoing change at each site was broadly categorized according to National Land Cover Dataset classes (Fry et al. 2011) as either desert (*n =* 12), forest (*n =* 40), grassland (*n =* 14), shrub/scrub (*n =* 10), or wetland (*n =* 14). Sites were chosen opportunistically while systematically balancing sample sizes among non-forested land cover types. At each study site, we manually delineated polygons in all areas of real change, and categorized observed changes as either bare ground, building (residential and commercial development), paved (roads, parking lots, etc.), or solar development. We then sampled the algorithm output values at all pixels within change area(s), and the maximum of either an equal number of random pixels or 1,000 random pixels within the study area not falling within areas of change and split these data into 70% training and 30% validation sets.

We used a combination of linear discriminant analysis and receiver operating characteristics to create sets of thresholds delineating changed and unchanged pixels based on algorithm outputs. First, we estimate the coefficients for a linear transformation of algorithm outputs (*CV, dNDVI*, etc. for LCC; *V1, V2*, …, *X*^*2*^ for MAD) into a single discriminant score that maximized the differentiation between algorithm outputs in changed and unchanged pixels. Coefficients were estimated from training data for changes occurring in all habitats, and specific to major habitat types. We then used receiver operating characteristic curves to identify the discriminant score providing greatest separation between true and false positives among validation data and assessed the performance of each algorithm in terms of the area under the curve. We identified the discriminant score that maximized the ratio between true and false positive rates as a threshold for automatically identifying changes. Linear discriminant and receiver operating characteristic analyses were conducted in R (Team 2014) using the *pscl* (Jackman 2017) and *pROC* (Robin et al. 2011) packages (Supporting Information).

### Case Studies

To demonstrate how these methods might be applied in situ, we evaluated the outputs from both change detection algorithms in each of four case studies (Table 1). These case studies were chosen as a sample of ongoing threats to imperiled species in a diversity of non-forested habitats. We focused outside of forested areas due to the extensive work and tools available for detecting deforestation (e.g., Global Forest Watch). Additionally, each case study represents a different potential use case for automated change detection; large-scale retrospective detection (dune sagebrush lizard); small-scale retrospective detection (beach mouse); rapid inventory after a natural disaster (Piping Plover); and active small-scale monitoring (Greater Sage Grouse).

**Table 1.** Study areas used in algorithm case studies covered a range of non-forested habitats and disturbances affecting imperiled species in the United States.

| *Species* | *Location* | *Habitat* | *Disturbance* | *Dates\** |
| --- | --- | --- | --- | --- |
| Greater Sage Grouse<br>(*Centrocercus urophasianus*) | Wright, WY | Grassland | Oil & gas | Jun., 2017 -<br>Sept., 2017 |
| Dune sagebrush lizard<br>(*Sceloporus arenicolus*) | Permian Basin,<br>NM & TX | Shrub/Scrub | Sand mining | Jan., 2017 -<br>Jan., 2018 |
| Beach mouse ssp.<br>(*Peromyscus polionotus*) | Gulf County,<br>FL | Wetland/<br>Grassland | Residential<br>construction | Jan., 2017 -<br>Jan., 2018 |
| Piping Plover<br>(*Charadrius melodus*) | Panama City,<br>FL | Grassland | Hurricane<br>Michael | Aug., 2018 -<br>Oct., 2018 |

\*Dates indicate the 'after' period during which changes were detected. A one-year interval preceding the earlier date was used as the 'before' period.

To evaluate algorithm effectiveness in these case studies, we compared algorithm outputs to changes identified by visual inspection of before and after images. Within each study area, an independent reviewer manually delineated all anthropogenic changes in the habitat type of interest. We refer to these polygons as ‘ground truth’ polygons. We then ran both the LCC and MAD algorithms within each study area, and delineated pixels representing change using the thresholds identified during LDA analysis. These areas representing change were then converted to polygons.

We use two complementary metrics to assess the algorithms’ performance. First, the Jaccard index measures the area of overlap between two geometries as the intersection divided by the union; *J*(*A,B*) = *A* ⋂ *B/A* ⋃ *B*. Second, we calculate the omission and commission error rates as the proportion of ground truth polygons that did not overlap algorithm output and the proportion of algorithm output that did not overlap any ground truth polygons, respectively.

In practice, we would apply a majority filter to these binary results to eliminate single, isolated pixels and create more contiguous areas of change or no change before conversion to polygons. However, to identify potential scale dependencies in algorithm performance, we converted to polygons all pixels identified as change. We then considered only sets of polygons greater than a sequence of minimum size {0 ac, 0.1 ac, 0.5 ac, 1 ac}, and calculate performance metrics within each of these subsets.

## Results

### Algorithm Validation

We collected algorithm output data from areas of real and no change at 100 locations (Supporting Information). Bare ground was the most common form of disturbance (86/100). Because bare ground preceded residential development and pavement, we did not detect these disturbance forms at any location. In 12 instances, solar fields were built directly over existing desert and grassland areas within the three-months comprising the ‘after’ image, and therefore appeared as a direct change from habitat to solar development.

Overall, both algorithms effectively discriminated change from no-change among validation data, as indicated by AUC scores > 0.90 for all habitat types (Fig. 2). Generally, the MAD algorithm performed slightly better than the LCC algorithm, as indicated by higher AUC scores. This was especially true in detecting ‘generic’ change, when thresholds were not optimized to a specific habitat type (Fig. 2). The LCC algorithm was least successful at identifying changes in grassland habitats (*AUC* = 0.95), and most successful in forested areas (*AUC* = 0.99). The MAD algorithm was most successful in wetland habitats (*AUC* = 1), and the least successful in forests (*AUC* = 0.98).

**Figure 2.**
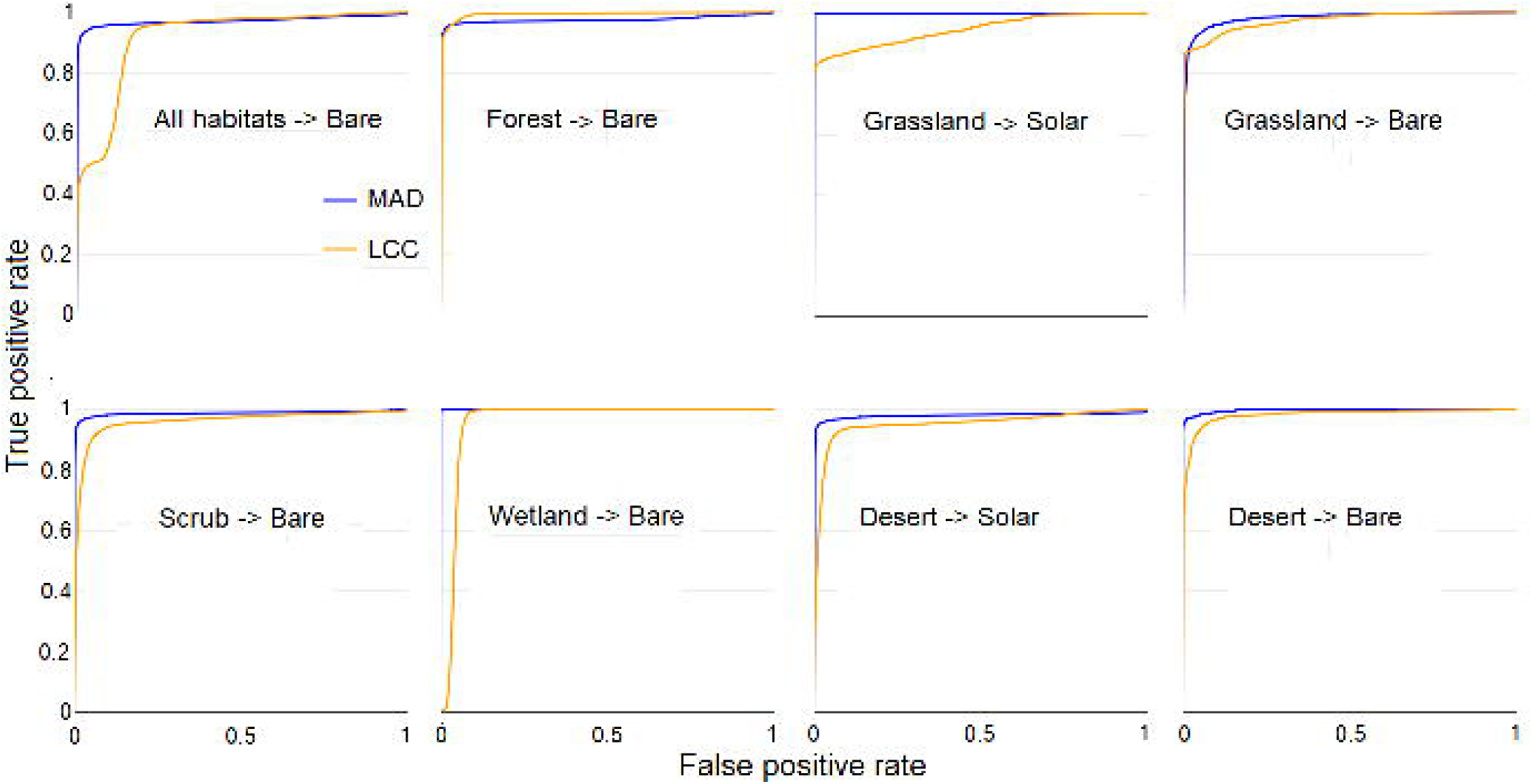
Receiver operating characteristic curves are used to identify thresholds for change detection. Curves plot the true and false positive rates for change detection among validation data as the linear discriminant analysis scores used as a delineating threshold increases. Curves constructed from the MAD algorithm outputs are shown in blue, and LCC algorithm outputs in orange. The values at which the rate of increase in detection rate relative to false positive rate decreases most rapidly are selected as threshold values. Curves are displayed for algorithm output data collected in different habitat types, and for all habitat types combined.

### Case Studies

We found these change detection methods were effective for detecting habitat loss in important conservation areas in all four case studies. We detected 8.5 km^2^ of dune sagebrush lizard habitat removed by either sand mining or oil and gas well construction within the Permian Basin, TX case study area (797.8 km^2^) between January 2017 and January 2018. The rapid appearance and expansion of large sand mines, in conjunction with ongoing oil and gas development identified by both algorithms (Fig. 3) indicated current protections for the imperiled lizard were insufficient to conserve the species.

**Figure 3.**
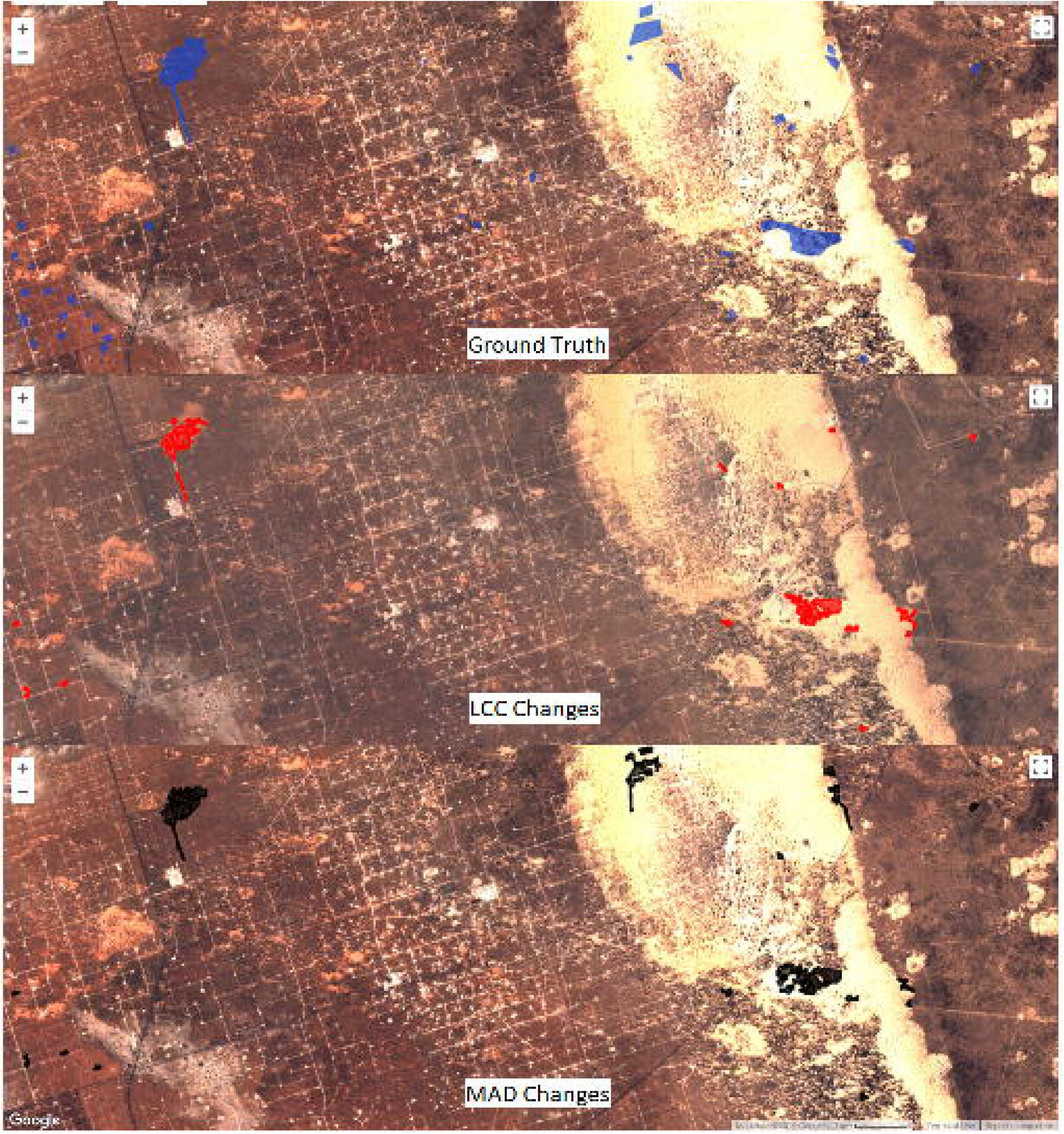
Habitat changes around a large dune complex in West Texas used by the dunes sagebrush lizard (*Sceloporus arenicolus*) were quickly identified by the LCC (red) and MAD (black) algorithms and align with those found by time-consuming manual delineation (blue). Changes were identified between pre-processed median composites from January 2018 as the before image, and January 2019 as the after image.

We detected 0.85 km^2^ of Piping Plover habitat within 26.2 km^2^ of designated critical habitat lost in the wake of Hurricane Michael (between August and October 2018), illustrating the threat posed by natural disasters to already imperiled species. In the Gulf County, FL case study area (4.7 km^2^), 0.03 km^2^ of potential beach mouse habitat were lost to residential development between January 2017 and January 2018. These areas of loss identified by automated change detection must be considered by the U.S. Fish and Wildlife Service when permitting future development in the species range. In the Wright, WY case study area (215.7 km^2^) we detected 0.17 km^2^ of grassland habitat loss between June and September 2017. This loss was due to oil and gas drilling pad and road construction.

The MAD algorithm was more sensitive to landscape changes, but less specific than the LCC algorithm, as indicated by higher commission rates and lower omission rates (Table 2). Both algorithms had relatively high commission rates, indicating detection of changes not identified by manual inspection of before and after images. A post-hoc analysis of commission indicated ∼60% of these polygons represented real changes missed by manual inspection. Jaccard indices indicated low to moderate agreement in the area of overlap between change polygons delineated manually and those produced by both automated change detection algorithms (Table 2). Commission rates decreased substantially when the minimum polygon size considered as change was increased from zero.

**Table 2.** Metrics of agreement between areas of change delineated by human review and automated change detection algorithms, using different minimum size thresholds for change polygons.

| Study Area | Min. Size (ac.) | MAD algorithm |  |  | LCC algorithm |  |  |
| --- | --- | --- | --- | --- | --- | --- | --- |
|  |  | *Jaccard*<sup>a</sup> | *Commission*<sup>b</sup> | *Omission*<sup>b</sup> | *Jaccard* | *Commission* | *Omission* |
| Wright, WY | 0.0 | 0.17 | 0.40 | 0.00 | 0.22 | 0.15 | 0.00 |
|  | 0.1 | 0.16 | 0.30 | 0.10 | 0.21 | 0.12 | 0.00 |
|  | 0.5 | 0.13 | 0.23 | 0.27 | 0.17 | 0.00 | 0.36 |
| Gulf County, FL | 0.0 | 0.41 | 0.67 | 0.04 | 0.46 | 0.40 | 0.14 |
|  | 0.1 | 0.44 | 0.32 | 0.28 | 0.47 | 0.10 | 0.23 |
|  | 0.5 | 0.31 | 0.00 | 0.80 | 0.33 | 0.00 | 0.80 |
| Panama City, FL | 0.0 | 0.23 | 0.41 | 0.11 | 0.26 | 0.33 | 0.15 |
|  | 0.1 | 0.25 | 0.27 | 0.19 | 0.26 | 0.17 | 0.22 |
|  | 0.5 | 0.22 | 0.13 | 0.28 | 0.24 | 0.05 | 0.31 |
| Permian Basin, TX | 0.0 | 0.36 | 0.89 | 0.04 | 0.37 | 0.79 | 0.08 |
|  | 0.1 | 0.37 | 0.72 | 0.11 | 0.37 | 0.58 | 0.15 |
|  | 0.5 | 0.39 | 0.39 | 0.34 | 0.37 | 0.29 | 0.43 |
|  | 1 | 0.39 | 0.17 | 0.42 | 0.36 | 0.13 | 0.60 |

<sup>a</sup>Jaccard index measures the degree of overlap between two sets of polygons on a zero (no overlap) to one (perfect overlap) scale
<sup>b</sup>Commission and omission rates were measured in terms of the number of polygons exclusive to either the ground truth (omission) or algorithm output (commission) sets.

Finally, algorithms detected changes between before and after images faster than human review. Both the LCC and MAD algorithms took < 40 min to produce change polygons in each study area. The time required for manual delineation of changes ranged from 6 hours in the Wright, WY case study area to several days in the Permian Basin, TX study area.

## Discussion

The conservation of biodiversity has been limited, in part, by an inability to monitor and enforce conservation laws, regulations, and agreements. While remote sensing data have long held the promise of transforming environmental monitoring efforts, publicly accessible tools leveraging this data to achieve actionable insights have been lacking (Willis 2015). In addition to cost, ease of use is critical if these tools are to be widely adopted for conservation monitoring and enforcement, as many land managers and regulators will not have expertise in ecology, policy, and remote sensing (Wiens et al. 2009). In this paper we adapt and present two algorithms for automated habitat change detection using satellite imagery, and demonstrate their efficacy, efficiency, and flexibility in a variety of test areas and case studies. Built on publicly available data and technology, these tools can be used by anyone - from local property managers to government agencies charged with national monitoring programs - to automatically detect habitat loss (Supporting Information).

Both the MAD (Nielsen 2007) and LCC (Jin et al. 2013) algorithms exhibited excellent performance discriminating habitat loss from background changes between images in test cases (Fig. 1). Beyond performing well in forested landscapes, where many algorithms have been developed (Hansen et al. 2010), both algorithms were effective in a variety of non-forest areas (Fig. 1). This flexibility is in part attributable to the use of land cover specific thresholds obtained from simple linear discriminant analysis using subsets of algorithm output data. We observed slightly lower AUC scores when receiver operating characteristic curves were produced using thresholds estimated from all data across land cover types. Specific thresholds were also important in detecting changes other than bare ground (e.g., solar energy development). The availability of a flexible tool that can be applied in a variety of contexts, rather than requiring a different tool for different ecosystems, should make automated change detection more readily adopted by entities with regulatory authority.

The ability of each algorithm to detect meaningful change was confirmed in case studies, where both methods identified nearly all instances of anthropogenic habitat loss that were manually delineated (i.e., low omission rates; Table 2). The MAD algorithm appeared more sensitive and less specific than LCC as illustrated by outputs from case studies. Generally higher commission rates among MAD outputs reflect the tendency of this algorithm to detect all types of change - even those occurring naturally due to phenology and seasonality. The change metrics included in the LCC algorithm that related to real-world phenomena (e.g., *dNDVI, dNBR*, etc.) likely enabled better discrimination between generic and habitat-specific change. Commission occurred from one of two outcomes: instances of habitat loss missed by manual review, or natural changes to the landscape that were not of interest. Object oriented, or computer vision-based approaches may be helpful for distinguishing among these (Malof et al. 2016; Ghorbanzadeh et al. 2019). Here, we present algorithms that are ready to be applied to a variety of habitats using only a Google Earth Engine account. Future work that integrates dynamically updated machine learning classification approaches, rather than a predefined set of thresholds, may also improve discrimination.

However, most instances of commission were not errors, illustrating a key advantage of automated change detection methods for conservation monitoring and enforcement: Both algorithms may be more effective than human review, particularly over large areas. The finding that ∼60% of instances of commission by both algorithms were in fact true cases of habitat loss demonstrates the potential for an automated change detection system to produce more complete result, especially over large areas, than manual inspection of before and after images. Furthermore, both algorithms were more efficient than manual inspection of satellite imagery. Human delineation of changes required several orders of magnitude more time to complete than automated algorithms and scaled with the size of the area of interest. Thus, automated change detection methods are vastly more time efficient, making them applicable and preferable in situations that require repeated or continuous monitoring. For entities wanting to use satellite imagery to implement a comprehensive conservation monitoring and enforcement program, this kind of efficiency is critical.

In the context of environmental laws that protect habitat, these automated change detection approaches could be used by regulatory agencies to enforce prohibitions on habitat destruction. In the United States, federal agencies responsible for implementing the Endangered species Act might use tools like this to monitor and enforce compliance with the terms and conditions of federal consultations under section 7 of the ESA, or habitat management plans associated with conservation agreements under section 10. These potential applications are illustrated by our case studies. For example, Gulf County, Florida has been developing a Habitat Conservation Plan to offset harm to endangered St. Andrew beach mice due to residential construction. These algorithms can be used to measure past and ongoing development and inform the plan as it is developed, as well as to monitor for compliance in the future. Additionally, the extent of historic habitat loss must be considered in future permitting decisions. Similarly, the ability to identify and track the expansion of sand mines within the range of the dune sagebrush lizard provided evidence that a state-run voluntary conservation agreement was insufficient to minimize threats faced by the species, and informed a petition to list the species under the ESA (https://ecos.fws.gov/docs/petitions/92210//1040.pdf). Finally, conservation agreements may often involve specifications of where development can and cannot occur within an area. The Wright, Wyoming case study provides a hypothetical example of how a small area could be monitored with relatively short (∼3 month) frequency to detect habitat destruction. Here, the changes detected here were legal, but in other instances might alert an enforcement agency to unauthorized activities.

Our results demonstrate the capability of both the MAD and LCC algorithms to automatically detect habitat loss, but those hoping to use these approaches should be aware of several caveats. First, both algorithms were more effective at identifying the occurrence of change than accurately delineating the extent of those changes. This was reflected by relatively low (< 0.5) Jaccard index scores. Change thresholds designed for specificity rather than sensitivity will invariably exclude some real changes, meaning the area of detected change will underestimate area changed. Additionally, the commission rates of both algorithms decreased as the minimum size of changes considered increased. This pattern suggests a minimum size of disturbance that can be regularly detected by these algorithms using Sentinel-2 data. If disturbances < 500 m^2^ need to be detected, users may experience a higher number of false positives.

While the algorithms presented here were run using Sentinel-2 data, they were written generically and can be applied to other passive remote sensing systems with the requisite bands and rigorous orthorectification and co-registration. For instance, Landsat 8, which provides global coverage of 30-meter resolution imagery every 16 days, contains analogous bands to Sentinel-2 as well as a thermal band measuring surface temperature allowing for more robust detection of clouds (Zhu et al. 2015). Our cloud detection and masking approaches are imperfect with Sentinel-2 imagery and may not perform as well in very cloudy areas. While habitat specific parameters help, clouds may still occasionally be flagged as change. Using these algorithms with Landsat data may be useful in cases where some resolution can be sacrificed for more robust cloud removal. Generic change detection algorithms using SAR data, which is invariant to cloud cover, may also prove useful in future development (Nielsen et al. 2017; Rüetschi et al. 2019).

By adapting two change detection algorithms and validating their efficacy across a variety of habitats, we have developed tools to help enforce conservation laws and agreements with remote sensing data (Supporting Information). The approaches described here do not require remote sensing expertise and can be used by local land managers, as well as federal agencies responsible for administering national and international laws. In addition, they are flexible, run much more quickly than manual delineation, and can be run repeatedly in many different contexts and at large spatial scales, making them suitable for the monitoring and enforcement of environmental laws. Most importantly, they are built using publicly available data and computing platforms. Many previous tools have been limited in their use in regulatory capacities because they are only available for a fee under pay-for-service structures. For remote sensing data to be used to improve conservation, it is critical that platforms like Google Earth Engine continue to provide open access. The continued improvement of automated change detection methods and adoption by regulatory authorities holds the potential to close a significant gap in the protection of biodiversity.

## Supporting Information

A public app running change detection is available at https://defendersofwildlifegis.users.earthengine.app/view/dowacd. All code used for image processing and change detection in Google Earth Engine, R code used to perform statistical analyses, and algorithm output data used for validation are available in an Open Science Framework repository (DOI: 10.17605/OSF.IO/5QAD8). The authors are solely responsible for the content and functionality of these materials. Queries (other than absence of the material) should be directed to the corresponding author.

## Acknowledgements

The authors would like to thank K. Stewart, M. Evansen, K. Blongewicz, and S. Steingard for their help with data collection. Thanks to J. Miller, M. Evansen, S. Steingard, and S. Patsel for reviewing the manuscript draft.

## Literature Cited

Betts MG, Wolf C, Ripple WJ, Phalan B, Millers KA, Duarte A, Butchart SHM, Taal L. 2017. Global forest loss disproportionately erodes biodiversity in intact landscapes. Nature 547:441–444.

Ceballos G, Ehrlich PR, Dirzo R. 2017. Biological annihilation via the ongoing sixth mass extinction signaled by vertebrate population losses and declines. Proceedings of the National Academy of Sciences 114:6089–6096.

Chandra A, Idrisova A. 2011. Convention on Biological Diversity: a review of national challenges and opportunities for implementation. Biodiversity and Conservation 20:3295–3316.

Chapron G. 2017. The environment needs cryptogovernance. Nature 545:403–405.

Drusch M et al. 2012. Sentinel-2: ESA’s Optical High-Resolution Mission for GMES Operational Services. Remote Sensing of Environment 120:25–36.

Farr TG et al. 2007. The Shuttle Radar Topography Mission. Reviews of Geophysics 45:RG2004.

Fry JA, Xian G, Jin S, Dewitz JA, Homer CG, Limin Y, Barnes CA, Herold ND, Wickham JD. 2011. Completion of the 2006 national land cover database for the conterminous United States. Photogrammetric Engineering and Remote Sensing 77:858–864.

Ghorbanzadeh O et al. 2019. Evaluation of Different Machine Learning Methods and Deep-Learning Convolutional Neural Networks for Landslide Detection. Remote Sensing 11:196.

Gorelick N, Hancher M, Dixon M, Ilyushchenko S, Thau D, Moore R. 2017. Google Earth Engine: Planetary-scale geospatial analysis for everyone. Remote Sensing of Environment 202:18–27.

Government Accountability Office. 2009. Endangered Species Act: The U.S. Fish and Wildlife Service has incomplete information about effects on listed species from section 7 consultations. Washington, DC.

Hansen MC, Stehman S V, Potapov P V. 2010. Quantification of global gross forest cover loss. Proceedings of the National Academy of Sciences of the United States of America 107:8650–5.

Hossain ANM, Barlow A, Barlow CG, Lynam AJ, Chakma S, Savini T. 2016. Assessing the efficacy of camera trapping as a tool for increasing detection rates of wildlife crime in tropical protected areas. Biological Conservation 201:314–319.

Hussey NE et al. 2015. Aquatic animal telemetry: A panoramic window into the underwater world. Science 348:1255642.

Jackman S. 2017. pscl: Classes and methods for R developed in the Political Science Computational laboratory. United States Studies Centre, University of Sydney, Sydney, New South Wales, AUstralia.

Japan Naitonal Diet. 1972. Nature Conservation Law.

Jin S, Yang L, Danielson P, Homer C, Fry J, Xian G. 2013. A comprehensive change detection method for updating the National Land Cover Database to circa 2011. Remote Sensing of Environment 132:159–175.

Keane A, Jones JPG, Edwards-Jones G, Milner-Gulland EJ. 2008. The sleeping policeman: Understanding issues of enforcement and compliance in conservation. Animal Conservation 11:75–82.

Lancaster HO, Seneta E. 2005. Chi-Square Distribution. Page Encyclopedia of Biostatistics. John Wiley & Sons, Ltd, Chichester, UK.

López-Bao JV et al. 2015. Toothless wildlife protection laws. Biodiversity and Conservation 24:2105–2108.

Malcom J. 2017. ESA Compliance Monitoring and the Langboard HCP. Washington, DC. https://defenders-cci.org/working_papers/Langboard_HCP/

Malcom J, Kim T, Li Y-W. 2017. Free Aerial Imagery as a Resource to Monitor Compliance with the Endangered Species Act. bioRxiv:204750.

Malcom JW, Li YW. 2015. Data contradict common perceptions about a controversial provision of the US Endangered Species Act. Proceedings of the National Academy of Sciences of the United States of America 112:15844–15849.

Malof JM, Bradbury K, Collins LM, Newell RG. 2016. Automatic detection of solar photovoltaic arrays in high resolution aerial imagery. Applied Energy 183:229–240.

McCarthy DP et al. 2012. Financial costs of meeting global biodiversity conservation targets: current spending and unmet needs. Science 338:946–949.

New Zealand Parliament. 1987. Conservation Act 1987.

Newbold T et al. 2015. Global effects of land use on local terrestrial biodiversity. Nature 520:45–50.

Nielsen AA. 2007. The regularized iteratively reweighted MAD method for change detection in multi- and hyperspectral data. IEEE Transactions on Image Processing 16:463–477.

Nielsen AA, Canty MJ, Skriver H, Conradsen K. 2017. Change detection in multi-temporal dual polarization Sentinel-1 data. Pages 3901–3904 IEEE International Geoscience and Remote Sensing Symposium. Fort Worth, TX.

Potapov P et al. 2008. Mapping the World’s Intact Forest Landscapes by Remote Sensing. Ecology and Society 13.

Robin X, Turck N, Hainard A, Tiberti N, Lisacek F, Sanchez J-C, Muller M. 2011. pROC: an open-source package for R and S+ to analyze and compare ROC curves. BMC Bioinformatics 12:77.

Rüetschi M, Small D, Waser L, Rüetschi M, Small D, Waser LT. 2019. Rapid Detection of Windthrows Using Sentinel-1 C-Band SAR Data. Remote Sensing 11:115.

Salomon M, Markus T, Dross M. 2014. Masterstroke or paper tiger – The reform of the EU’s Common Fisheries Policy. Marine Policy 47:76–84.

Sarabandi P, Yamazaki F, Matsuoka M, Kiremidjian A. 2004. Shadow detection and radiometric restoration in satellite high resolution images. Pages 3744–3747 International Geoscience and Remote Sensing Symposium (IGARSS).

Song X-P, Hansen MC, Stehman S V., Potapov P V., Tyukavina A, Vermote EF, Townshend JR. 2018. Global land change from 1982 to 2016. Nature 560:639–643.

R Core Development Team. 2019. R: a languange and environmental for statistical computing. Version 3.5.1. R Foundation for Statistical Computing, Vienna, Austria.

Teillet PM, Guindon B, Goodenough DG. 1982. On the Slope-Aspect Correction of Multispectral Scanner Data. Canadian Journal of Remote Sensing 8:84–106.

Trouwborst A et al. 2017. International Wildlife Law: Understanding and Enhancing Its Role in Conservation. BioScience 67:784–790.

Turner W, Spector S, Gardiner N, Fladeland M, Sterling E, Steininger M. 2003. Remote sensing for biodiversity science and conservation. TRENDS in Ecology and Evolution 18:306–314.

UN Environment World Conservation Monitoring Centre, International Union for Conservation of Nature. 2017. World Database on Protected Areas. Available from http://www.protectedplanet.net

United States Congress. 1978. The Endangered Species Act Amendments of 1978.

Waldron A, Mooers AO, Miller DC, Nibbelink N, Redding D, Kuhn TS, Timmons Roberts J, Gittleman JL. 2013. Targeting global conservation funding to limit immediate biodiversity declines. Proceedings of the National Academy of Science 110:12144–12148.

Wiens JA, Stralberg D, Jongsomjit D, Howell CA, Snyder MA. 2009. Niches, models, and climate change: assessing the assumptions and uncertainties. Proceedings of the National Academy of Sciences of the United States of America 106:19729–36.

Willis KS. 2015. Remote sensing change detection for ecological monitoring in United States protected areas. Biological Conservation 182. 233–244.

Xu H. 2006. Modification of normalised difference water index (NDWI) to enhance open water features in remotely sensed imagery. International Journal of Remote Sensing 27:3025–3033.

Zhu Z, Wang S, Woodcock CE. 2015. Improvement and expansion of the Fmask algorithm: Cloud, cloud shadow, and snow detection for Landsats 4-7, 8, and Sentinel 2 images. Remote Sensing of Environment 159:266–277.

Zhu Z, Woodcock CE. 2012. Object-based cloud and cloud shadow detection in Landsat imagery. Remote Sensing of Environment:83–94.

